# Heartbeat perception is causally linked to frontal delta oscillations

**DOI:** 10.1101/2024.05.25.595873

**Authors:** David Haslacher, Philipp Reber, Alessia Cavallo, Annika Rosenthal, Elisabeth Pangratz, Anne Beck, Nina Romanczuk-Seiferth, Vadim Nikulin, Arno Villringer, Surjo R. Soekadar

## Abstract

The ability to accurately perceive one’s own bodily signals, such as the heartbeat, plays a vital role in physical and mental health. However, the neurophysiological mechanisms underlying this ability, termed interoception, are not fully understood. Converging evidence suggests that cardiac rhythms are linked to frontal brain activity, particularly oscillations in the delta (0.5 – 4 Hz) band, but their causal relationship remained elusive. Using amplitude-modulated transcranial alternating current stimulation (AM-tACS), a method to enhance or suppress brain oscillations in a phase-specific manner, we investigated whether frontal delta oscillations are causally linked to heartbeat perception. We found that enhancement of delta phase synchrony suppressed heartbeat detection accuracy, while suppression of delta phase synchrony enhanced heartbeat detection accuracy. These findings suggest that frontal delta oscillations play a critical role in heartbeat perception, paving the way for causal investigations of interoception and potential clinical applications.

**Significance:** Although bodily signals are known to influence perception and behavior, little is known about the underlying neurophysiological mechanisms. Here, we show that perception of the heartbeat is anticorrelated with phase synchrony of frontal delta oscillations, and that modulating these oscillations with transcranial electric stimulation influences heartbeat perception. Our results suggest that delta oscillations play a key role in processing bodily signals, with potential implications for theories of emotions and clinical neuroscience.

## Introduction

Interoception refers to the sense by which individuals perceive the physiological state of their body, such as satiation, hydration, respiration, and the heartbeat (1). It was found that accurate interoception is essential for the effective regulation of physiological processes, ensuring that the body’s response to environmental challenges is appropriate. For instance, accurate perception of hydration leads to appropriate fluid intake, which in turn maintains body function (2). Moreover, the integrity of interoception ensures that cardiovascular, metabolic, and other physiological states can be adjusted to the individual’s current and future requirements (3, 4). In turn, dysfunctions in interoception were found in a range of mental health conditions including anxiety (5), depression (6), panic disorder (7), eating disorders (8), and substance use disorders (9).

Similarly to sensorimotor processes, interoception forms a perception-action loop (10). Interoceptive signals, originating from sensory receptors within internal organs, reflect information about physiological states that is integrated with other information in the insular and anterior cingulate cortices (1, 4, 11). Efferent signals are then generated in the form of behavioral responses (e.g., eating or drinking) or autonomic adjustments (e.g., increasing heart rate). This bidirectional interplay ensures an adaptive response to both perturbations of the body state and environmental demands (10, 12).

It was found that the interaction between brain and body is intrinsically rhythmic, spanning over a wide range of frequencies (13). For instance, the gastric basal rhythm oscillates at about 0.05 Hz, respiration at about 0.25 Hz, and the heartbeat at about 1 Hz. Although the link between these rhythms and those of the brain is unclear, there is increasing evidence of both afferent and efferent forms of temporal locking in the sense of *entrainment* (14-19). Moreover, several studies found various links between frontal delta oscillations (FDOs, 0.5 – 4 Hz) and autonomic functions (20), such as an anticorrelation between frontal delta power and the heart rate (18). At the same time, it was found that cardiac activity can modulate FDOs (19, 21). These results suggest that FDOs play an important role in the bidirectional coupling between the brain and the heart. However, it is not clear how this coupling relates to heartbeat perception. Since brain responses related to self-generated sensory input are typically suppressed (22-24), we hypothesized that phase synchrony of FDOs are causally and negatively linked to heartbeat-evoked potentials (HEPs) and heartbeat perception.

To test this hypothesis, we invited healthy human volunteers (N = 24) to perform an established heartbeat detection task (11) while electroencephalography (EEG) was recorded (Fig. 1). In this task, participants had to indicate whether an auditory tone sequence was presented early or late relative to the heartbeat. To differentiate EEG signals related to auditory processing, participants also performed an auditory tone detection task in which they were asked to indicate deviant tones within a tone sequence.

**Figure 1.**
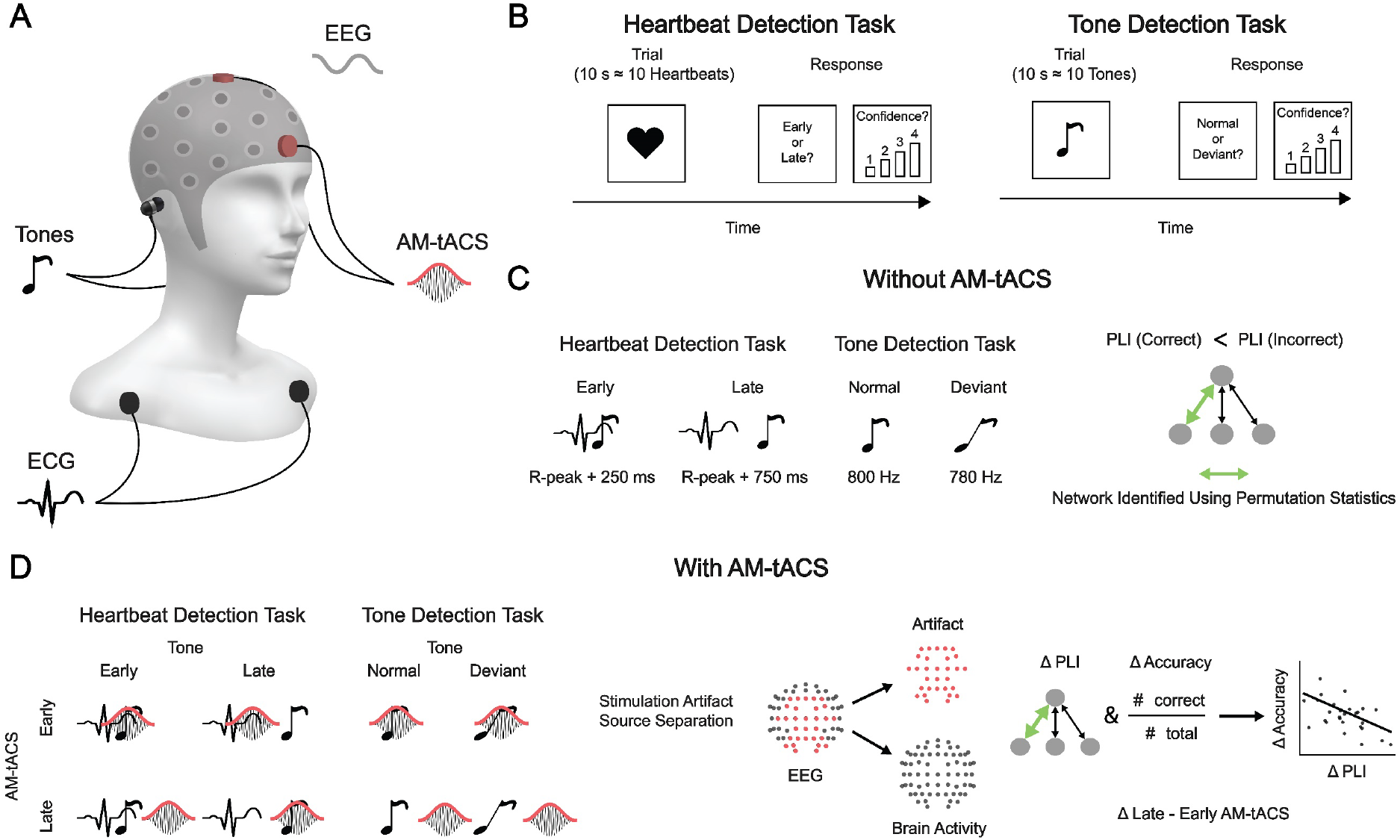
Experimental paradigm. **(A)** Electroencephalography (EEG) and electrocardiography (ECG) were recorded while auditory stimuli were delivered through a pair of earphones. To modulate delta phase synchrony, amplitude-modulated transcranial alternating current stimulation (AM-tACS) was applied to the frontal cortex. Parameters of AM-tACS comprised an 8 kHz carrier frequency, an envelope frequency corresponding to the heart rate, and a stimulation intensity adjusted to avoid any somatosensory perception, up to ± 10 mA. **(B)** During the heartbeat detection task (11), participants (N = 24) were asked to assess whether their heartbeat was early or late relative to the sequence of pure tones. During the tone detection task, participants were asked to assess whether the sequence of pure tones contained a deviant tone. Auditory stimuli were identical across tasks, and trials of each task were mixed such that they were performed in a pseudorandomized order. **(C)** In absence of AM-tACS, pure tones were played either early with the perceived heartbeat, (∼250 ms after ECG R-peak) or late (∼750 ms after ECG R-peak). Auditory stimuli consisted of a sequence of normal (800 Hz) tones, in some trials containing a deviant (780 Hz) tone. To identify the network of delta oscillations linked to heartbeat perception, a network-based permutation test was used to assess where the phase lag index (PLI) was higher during incorrect than during correct responses in the heartbeat detection task. **(D)** Participants performed the same tasks in the presence of AM-tACS, which was applied early or late relative to (perception of) the heartbeat. We show only one cycle of AM-tACS for illustrative purposes, although it was applied (and adjusted to the heartbeat) continuously. Stimulation artifact source separation (SASS) was used for stimulation artifact removal from EEG signals. Finally, modulation of delta phase synchrony in the network previously identified was correlated with the modulation of heartbeat detection accuracy.

Assessing the entrainment of brain oscillations by interoceptive signals such as the heartbeat comes with methodological challenges. By aligning their high- or low-excitability phases with rhythmic input, brain oscillations constitute an efficient mechanism for filtering of predictable sensory signals (25, 26). Typically, this phenomenon is studied by comparing the phase of brain oscillations at the time of enhanced and suppressed sensory input (26, 27). However, due to heartbeat-locked artifacts in the EEG (28-30), the phase of delta oscillations relative to the heartbeat is difficult to assess directly. Instead, since coordinated entrainment of brain oscillations manifests in network synchrony (31), we chose to assess delta phase synchrony between brain regions by computing the phase lag index (PLI) to mitigate the influence of cardiac artifacts (32). Subsequently, we applied amplitude-modulated transcranial alternating current stimulation (AM-tACS), a frequency-tuned form of non-invasive brain stimulation to enhance or suppress brain oscillations (33), to the frontal cortex either early or late relative to (perception of) the heartbeat. We expected that AM-tACS results in phase-dependent enhancement and suppression of delta phase synchrony in the identified network causing an increase and decrease in heartbeat detection accuracy.

## Results

### Frontal delta phase synchrony is associated with heartbeat perception

We found that delta phase synchrony was associated with heartbeat detection accuracy in a network of frontal brain regions (Fig. 2A), which exhibited higher PLI during incorrect than during correct responses (t(23) = 1.61, p = 0.044, network-based permutation test), confirming that delta phase synchrony was negatively associated with heartbeat detection accuracy. We also assessed whether PLI in the identified network was larger during the tone detection task than during the heartbeat detection task (Fig. 2B). No such difference was found (t(23) = 0.281, p = 0.623, permutation test), demonstrating that increased delta phase synchrony was associated with decreased heartbeat perception but not tone perception. Correspondingly, a two-way ANOVA revealed an interaction between task (heartbeat or tone detection) and response (correct or incorrect) with respect to frontal delta synchrony (F(1,23) = 5.90, p = 0.0233). We also assessed whether differences in oscillatory synchrony between incorrect and correct responses were restricted to the delta band by performing the network-based permutation test for the theta (4 – 8 Hz), alpha (8 – 15 Hz), beta (15 – 30 Hz), and gamma (30 – 45 Hz) bands. No differences in synchrony between incorrect and correct responses were found in these frequency bands. Cluster-based permutation testing did not reveal any difference in delta amplitude between correct and incorrect responses.

**Figure 2.**
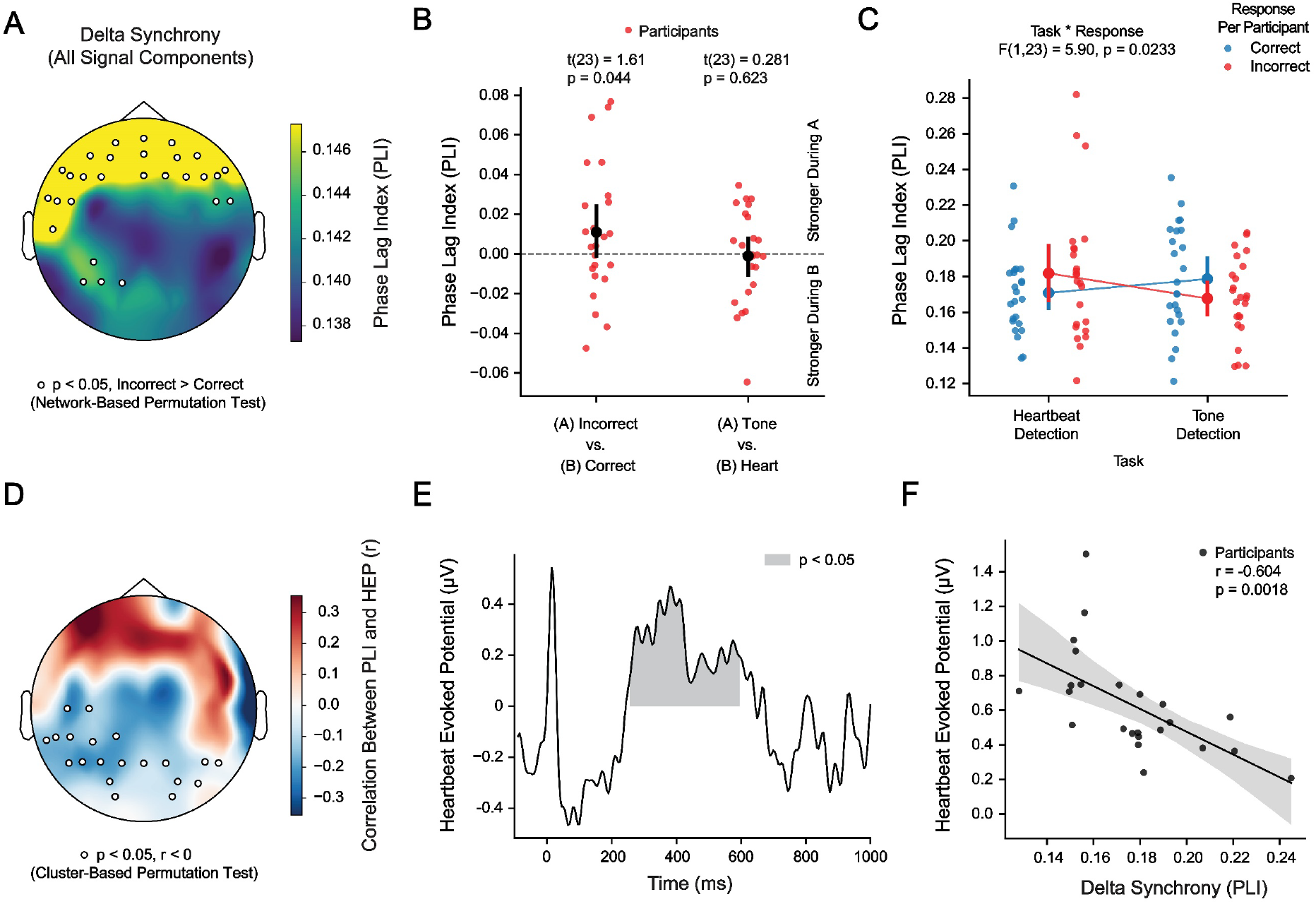
Increased frontal delta phase synchrony is associated with reduced heartbeat perception and reduced heartbeat-evoked potential. **(A)** Phase synchrony of delta oscillations recorded over frontal brain areas was associated with heartbeat detection. **(B)** Delta phase synchrony was larger during incorrect than during correct responses in the heartbeat detection task. No difference in delta phase synchrony between the heartbeat and tone detection tasks was found. **(C)** An interaction between task (heartbeat or tone detection) and response (correct or incorrect) was found with respect to frontal delta synchrony. **(D)** Delta phase synchrony was anticorrelated with the amplitude of the heartbeat-evoked potential (HEP) in several clusters of central, parietal, and occipital EEG electrodes. **(E)** The anticorrelation between delta phase synchrony and HEP amplitude was found between 255 and 595 ms after the ECG R-peak. **(F)** When averaged over significant electrodes and timepoints, delta phase synchrony was strongly anticorrelated with the HEP amplitude.

We next assessed whether PLI in the identified network was associated with HEP amplitude in any brain region. We found an anticorrelation between PLI (calculated over the entire trial duration) and HEP amplitude in central, parietal, and occipital areas from 255 to 595 ms after the ECG R-peak (p < 0.05, cluster-based permutation test) (Fig. 2D and 2E). The strength of this anticorrelation strongly increased when HEP amplitudes were averaged across significant sensors and timepoints (r = -0.604, p = 0.0018) (Fig. 2F). To further confirm that delta phase synchrony was associated with heartbeat perception and not tone perception, we assessed whether PLI in the identified network was associated with auditory evoked potential (AEP) amplitude. Cluster-based permutation testing did not reveal any correlation between PLI and AEP amplitude.

We also assessed whether evoked responses in the delta frequency range activity might explain our findings. In addition to endogenous frontal delta oscillations, heartbeat-evoked activity, as well as auditory-evoked activity, were observable during our task. We found that frontal delta synchrony, particularly its difference between incorrect and correct responses, was not accounted for by evoked responses. Neither was it accounted for by ocular artifacts (Fig. S1).

### Transcranial alternating current stimulation modulates frontal delta phase synchrony and heartbeat perception

We first assessed whether the timing of AM-tACS relative to the ECG R-peak influenced heartbeat perception (Fig. 3A). We found that heartbeat detection accuracy during late AM-tACS (66.0 ± 14.7 %) was lower (t(24) = -2.06, p = 0.0256) than during early AM-tACS (69.2 ± 15.7 %). Before assessing delta oscillations recorded in the presence of AM-tACS, we confirmed that electric stimulation artifacts were successfully attenuated by SASS (Fig. S1). We found that the change in delta phase synchrony across delay conditions predicted (r = - 0.411, p = 0.0231) the change in heartbeat detection accuracy (Fig. 3B). Thus, an increase in delta phase synchrony led to a decrease in heartbeat detection accuracy, and vice-versa. Cluster-based permutation testing did not reveal any difference in delta amplitude between AM-tACS delay conditions.

**Figure 3.**
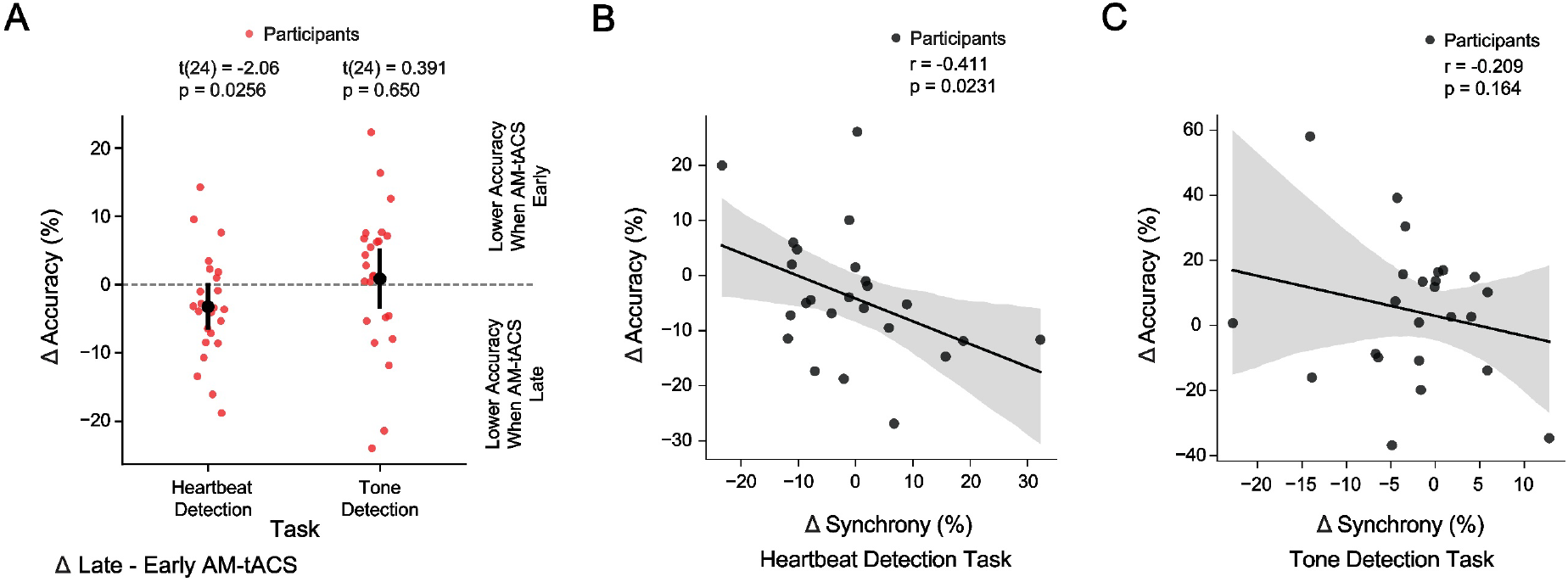
Amplitude-modulated transcranial alternating current stimulation (AM-tACS) suppresses heartbeat detection accuracy by enhancing frontal delta phase synchrony. **(A)** When AM-tACS was applied out of phase with the heartbeat, heartbeat detection accuracy was lower than when AM-tACS was applied early relative to the heartbeat. No differential effect on tone detection accuracy was found. **(B)** The modulation of delta phase synchrony in the target network (Fig. 2) was anticorrelated with the modulation of heartbeat detection accuracy, such that an enhancement of delta phase synchrony caused a suppression of heartbeat detection accuracy. **(C)** No relationship between the modulation of delta phase synchrony and tone detection accuracy was found.

To exclude that physiological and behavioral effects of AM-tACS are related to neural processing of the auditory stimulus, we performed the same analyses for the tone detection task. We found that tone detection accuracy during out of phase AM-tACS (49.7 ± 5.51 %) was comparable to (t(24) = 0.391, p = 0.650) tone detection accuracy during early AM-tACS (48.9 ± 7.11 %). No relationship (r = -0.209, p = 0.164) between the change in synchrony and the change in accuracy across delay conditions was found (Fig. 3C).

While we find a link between changes in frontal delta synchrony and changes in heartbeat detection accuracy due to AM-tACS (Fig. 3B), we did not find a difference in frontal delta synchrony between late and early AM-tACS conditions. We reasoned that this might result from a variability in the optimal phase difference between AM-tACS and frontal delta oscillations to enhance (or suppress) frontal delta synchrony. Indeed, we find that this optimal phase difference varies across participants (Fig. S4). We also find that this phase difference varies across participants within both the early and late AM-tACS conditions (Fig. S3).

## Discussion

Our findings indicate that FDOs suppress heartbeat-evoked brain activity, attenuating perception of the heartbeat. While previous studies have found that processing of visual (17, 34-36), auditory (37), and somatosensory (38-41) stimuli depends on the timing of stimulus presentation relative to the heartbeat, the underlying mechanism remained unknown. Our findings suggest that such heart cycle dependent modulations of perception may be mediated by FDOs, consistent with the notion that they might reflect unconscious predictions of bodily signals such as the heartbeat (40, 42-44). Importantly, our findings suggest that this theoretical concept is amenable to experimental investigation using phase-specific brain stimulation. Future studies should also investigate whether our findings can be generalized to the processing of other interoceptive signals beyond heartbeat (20), and how manipulation of these interoceptive signals affects FDOs (45).

Causal modulation of FDOs via AM-tACS offers an approach to verify their functional relevance for heartbeat perception and could be applied to experimentally test basic theories of emotion, which make competing predictions about the causal role of interoception. The James-Lange theory posits that we feel emotions in response to bodily signals (46). In contrast, the Cannon-Bard theory argues that bodily signals and emotional experiences occur simultaneously but independently (47). Meanwhile, the Schachter-Singer theory suggests that emotions arise from a combination of bodily signals and cognitive appraisal (48). By selectively manipulating the processing of bodily signals using AM-tACS, it might be possible to experimentally test these competing basic theories.

Finally, our results suggest that AM-tACS could be used to better understand whether alterations in interoception are causally related to symptoms of a range of health conditions. For instance, it has been proposed that altered interoceptive processing is a primary cause of anxiety and depression (5, 6), in addition to other psychiatric disorders (49). Dysfunctions of interoception have also been linked to a range of other brain disorders with somatic symptoms such as chronic pain (50), obesity (51), and chronic stress (52). Beyond disentangling the causal relationship between interoception and other aspects of these disorders, our approach might facilitate new avenues of treatment.

However, several methodological limitations of tACS should be addressed to reduce variability of stimulation effects across clinical applications (53-56). Effect variability may result from inter-individual anatomical differences, which should be considered in the design of personalized stimulation montages (55). Importantly, effects of tACS also depend on the brain state during stimulation (57-59). We found that AM-tACS enhanced and suppressed FDOs in a phase-dependent manner. While this mechanism has been documented elsewhere for other brain oscillations (33, 60-63), its utilization in clinical practice remains a challenge. To selectively enhance or suppress a brain oscillation using this mechanism, electric stimulation artifacts in simultaneously recorded EEG must be sufficiently attenuated to extract single-trial phase information and continually adapt tACS to the oscillation in real-time (33, 64, 65). Furthermore, the phase difference between tACS and the oscillation must be personalized to achieve the desired enhancement or suppression effect (33, 61, 63, 66). Finally, poor focality and depth of transcranial electric stimulation (54, 67, 68) limit the capacity of tACS to selectively modulate subregions of the frontal brain mediating interoception. Recently developed techniques employing interfering electric (69-71) or magnetic (72) fields may alleviate these issues while remaining compatible with the stimulation artifact rejection approaches employed here (33, 64, 65). These technical advances may enable more refined investigations of the causal link between FDOs, interoception, and brain (dys)function.

## Methods

### Participants

In total, 25 participants (14 female, 11 male, 26 ± 5 years of age) were invited to participate in the study and provided written informed consent in accordance with the ethics committee of the Charité – Universitätsmedizin Berlin (EA1/077/18). One participant was excluded due to exceedingly high heartbeat detection accuracy (> 90%) in all conditions.

### Electroencephalography

Electroencephalograph (EEG) was recorded from the scalp according to the international 10 – 20 system using a NeurOne system (Bittium Corp., Oulu, Finland). For all recordings, the amplifier was set to DC-mode with a dynamic range of +/-430 mV, a resolution of 51 nV/bit, and a range of 24 bit. Data were sampled at 2 kHz. The impedance for all sensors was kept below 10 kΩ. Otherwise, the sensors were marked for interpolation.

### Electrocardiography

Electrocardiography (ECG) was recorded from an auxiliary bipolar channel along with EEG data. Electrodes were placed under the left and right infraclavicular fossa. The EEG system sent the ECG data to a real-time computer via a real-time UDP stream, where it was further processed to control AM-tACS and auditory stimuli.

### Transcranial alternating current stimulation

Amplitude-modulated transcranial alternating current stimulation (AM-tACS) was delivered to the scalp with a current of up to ± 10 mA and a carrier signal frequency of 8 kHz using a Digitimer DS5 (Digitimer Ltd, UK). The amplitude of AM-tACS was adjusted individually for each participant to the maximal level that remained below the threshold for somatosensory perception. Two circular rubber electrodes (34 mm diameter, 2mm thickness) were used with conductive ten20 paste (Weaver & Co, Aurora, CO, USA) to apply AM-tACS to the scalp. Rubber electrodes delivering AM-tACS were centered on positions Fpz and Cz of the international 10-20 system. The electric stimulator was controlled by an SDG 2042X signal generator (Siglent, NL), which applied amplitude-modulation to the carrier signal depending on an input voltage signal received from the Speedgoat Performance Real-Time Target Machine (Speedgoat GmbH, CH). The real-time target machine adjusted the phase of amplitude envelope to the ECG signal in real-time, such that its maximum was either early (∼250 ms after ECG R-peak) or late (∼750 ms after ECG R-peak) with (perception of) the heartbeat.

### Auditory stimuli

Auditory stimulation consisted of a sequence of pure tones of 100 ms duration delivered through earphones at a 60 dB level. Tones were played either early (∼250 ms after ECG R-peak) or late (∼750 ms after ECG R-peak) relative to the heartbeat. The timing of early and late tones was chosen to match that of a prior study implementing the same paradigm, but with pulse measurements from the finger (11). In that study, early tones were played < 150 ms after the pulse wave, whereas late tones were played ∼500 ms after the pulse wave. Our timing was subsequently chosen based on evidence that the pulse wave arrives at the finger ∼250 ms after the ECG R-peak (73). Normal tones had a frequency of 800 Hz, whereas deviant tones had a frequency of 785 Hz.

### Behavioral tasks

Participants carried out randomized trials of a heartbeat detection task and a tone detection task. For sensory stimulation parameters, see the previous section. Throughout each 10-second trial of either task, they were presented with a series of tones, which was triggered by their own heartbeat. In 50% of trials, the tones were played early relative to the perceived heartbeat (∼250 ms after ECG R-peak). In the remaining 50%, the tones were played late relative to the perceived heartbeat (∼750 ms after ECG R-peak). Additionally, in 50% of the trials, the tone sequence contained a deviant tone. In the heartbeat detection task, we manipulated attention by presenting study participants with a heart symbol in the middle of the screen. At the end of each trial, they were asked to indicate whether the tones were early or late relative to the heartbeat, and how confident they were about their response (by selecting 1,2,3, or 4). Throughout the tone detection task, we directed them to attend to the pitch of the tones by showing them a musical note symbol in the middle of the screen. At the end of each trial, they were asked to indicate whether the tone sequence contained a deviant tone, and how confident they were about their response (by selecting 1,2,3, or 4). To compute heartbeat and tone detection accuracy, responses were weighted by their confidence.

### Stimulation artifact source separation

Following (64), we removed the AM-tACS artifact from bandpass-filtered (0.5 – 4 Hz) EEG data using SASS. First, we computed the sensor covariance matrix **A** in the presence of AM-tACS, as well as the sensor covariance matrix **B** in absence of AM-tACS. Subsequently, a source separation matrix W was computed by joint diagonalization of **A** and 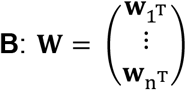 where 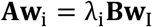 and 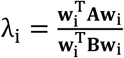. The **w**_i_ were ordered from greatest to least **λ**_I_, which represented the ratio of artifact power to brain signal power (noise-to-signal ratio) in each component. Finally, the matrix P was constructed as **P** = **W**^−1^**SW**, where 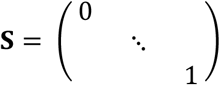, with zeros on the diagonal representing artifact components that were removed from the data. To remove AM-tACS artifacts, the sensor-space data were then multiplied by P. To select the number of components to reject (zeros in S), |**B** − **PBP**^**T**^|_2_ was minimized, resulting in 20.0 ± 11.1 rejected components.

### Electroencephalography data processing

To compute the HEP in absence of AM-tACS, EEG data were filtered from 0.5 - 30 Hz using a finite impulse response filter. Epochs were then extracted around the ECG R-peak, and the signal was averaged over epochs. To compute delta phase, EEG data were filtered from 0.5 – 4 Hz using a finite impulse response filter of order 1321 (corresponding to 6.61 seconds). For data recorded in the presence of AM-tACS, SASS was then applied to suppress the electric stimulation artifact (see previous section). Subsequently, the Hilbert transform was applied to the data, and the angle of the analytic signal was taken to obtain the phase of delta activity. To assess long-range delta phase synchrony, the phase lag index was computed between each pair of channels (32). MNE-Python (74), NumPy (75), and SciPy (76) were used for all analyses. To remove delta-band activity linked to heartbeat-evoked potentials (HEP) and auditory-evoked potentials (AEP) (Fig. S1), a template subtraction procedure was employed. First, the HEP and AEP were computed from – 500 ms to 1000 ms around the ECG R-peak and tone onset, respectively. Then, the template HEP and AEP were subtracted from EEG signals at each instance of the ECG R-peak and tone onset, respectively. To remove delta-band activity linked to ocular artifacts, a regression-based approach was employed (77). First, a virtual EOG channel was computed as the difference between signals from EEG electrodes Fp1 and Fp2. Then, linear regression was used to remove the resulting EOG signal from EEG signals.

### Statistics

To identify the network of delta oscillations whose synchrony differed across correct and incorrect responses in the heartbeat detection task, network-based statistics were employed (78). First, a threshold was computed as the 95^th^ percentile of all entries in the connectivity (PLI) matrix averaged across epochs and participants. Second, the largest connected component (network) was identified from the averaged and thresholded connectivity matrix. To assess statistical significance of synchrony in this network, the PLI was averaged within the network for each participant and condition (correct or incorrect responses in the heartbeat detection task). From this, a t-statistic was computed for the network. A permutation test was then used to compute a p-value for the network by shuffling the condition labels across trials within each participant and recomputing the t-statistic 10,000 times. To identify clusters across time and space where HEP amplitude correlated with delta phase synchrony in the identified network, a cluster-based permutation test was used (79). To obtain a test statistic, Pearson’s r was computed across participants between PLI in the network and HEP amplitudes at each point in time and space. In each permutation, PLI in the network was shuffled across participants. To assess opposition of the phase difference between AM-tACS and frontal delta oscillations across early and late AM-tACS conditions (Fig. S3), we employed the phase opposition sum (80). To assess phase-dependent modulation of frontal delta synchrony by AM-tACS (Fig. S4), we fit a sine function to the single-trial data using non-linear least-squares optimization. To assess significance of the phase opposition sum or amplitude of phase-dependent modulation within each participant, we employed a permutation test. The labels (condition or AM-tACS-delta phase difference) were permuted across trials, and the test statistic was recomputed 10,000 times.

## Acknowledgements

This project was supported by the Max Planck Institute Human Cognitive and Brain Sciences through the MBB Young Scientist Award given to David Haslacher. This work was also supported in part by the European Research Council (ERC) under the project NGBMI (759370) and TIMS (101081905), and the Einstein Stiftung Berlin.

## References

1. A. D. Craig, How do you feel? Interoception: the sense of the physiological condition of the body. Nat Rev Neurosci 3, 655–666 (2002).

2. R. J. Stevenson, M. Mahmut, K. Rooney, Individual differences in the interoceptive states of hunger, fullness and thirst. Appetite 95, 44–57 (2015).

3. B. S. McEwen, J. C. Wingfield, The concept of allostasis in biology and biomedicine. Horm Behav 43, 2–15 (2003).

4. I. R. Kleckner et al., Evidence for a Large-Scale Brain System Supporting Allostasis and Interoception in Humans. Nat Hum Behav 1 (2017).

5. M. P. Paulus, M. B. Stein, An insular view of anxiety. Biol Psychiatry 60, 383–387 (2006).

6. M. P. Paulus, M. B. Stein, Interoception in anxiety and depression. Brain Struct Funct 214, 451–463 (2010).

7. M. B. Stein, G. J. Asmundson, Autonomic function in panic disorder: cardiorespiratory and plasma catecholamine responsivity to multiple challenges of the autonomic nervous system. Biol Psychiatry 36, 548–558 (1994).

8. L. A. Berner et al., Altered interoceptive activation before, during, and after aversive breathing load in women remitted from anorexia nervosa. Psychol Med 48, 142–154 (2018).

9. R. Z. Goldstein et al., The neurocircuitry of impaired insight in drug addiction. Trends Cogn Sci 13, 372–380 (2009).

10. F. H. Petzschner, S. N. Garfinkel, M. P. Paulus, C. Koch, S. S. Khalsa, Computational Models of Interoception and Body Regulation. Trends Neurosci 44, 63–76 (2021).

11. H. D. Critchley, S. Wiens, P. Rotshtein, A. Ohman, R. J. Dolan, Neural systems supporting interoceptive awareness. Nat Neurosci 7, 189–195 (2004).

12. D. S. Ramsay, S. C. Woods, Clarifying the roles of homeostasis and allostasis in physiological regulation. Psychol Rev 121, 225–247 (2014).

13. W. Klimesch, The frequency architecture of brain and brain body oscillations: an analysis. Eur J Neurosci 48, 2431–2453 (2018).

14. W. Zhong et al., Selective entrainment of gamma subbands by different slow network oscillations. Proc Natl Acad Sci U S A 114, 4519–4524 (2017).

15. J. Ito et al., Whisker barrel cortex delta oscillations and gamma power in the awake mouse are linked to respiration. Nat Commun 5, 3572 (2014).

16. C. Zelano et al., Nasal Respiration Entrains Human Limbic Oscillations and Modulates Cognitive Function. J Neurosci 36, 12448–12467 (2016).

17. H. D. Park, S. Correia, A. Ducorps, C. Tallon-Baudry, Spontaneous fluctuations in neural responses to heartbeats predict visual detection. Nat Neurosci 17, 612–618 (2014).

18. E. Patron, R. Mennella, S. Messerotti Benvenuti, J. F. Thayer, The frontal cortex is a heart-brake: Reduction in delta oscillations is associated with heart rate deceleration. Neuroimage 188, 403–410 (2019).

19. D. Candia-Rivera, V. Catrambone, J. F. Thayer, C. Gentili, G. Valenza, Cardiac sympathetic-vagal activity initiates a functional brain-body response to emotional arousal. Proc Natl Acad Sci U S A 119, e2119599119 (2022).

20. G. G. Knyazev, EEG delta oscillations as a correlate of basic homeostatic and motivational processes. Neurosci Biobehav Rev 36, 677–695 (2012).

21. D. Candia-Rivera, V. Catrambone, R. Barbieri, G. Valenza, Functional assessment of bidirectional cortical and peripheral neural control on heartbeat dynamics: A brain-heart study on thermal stress. Neuroimage 251, 119023 (2022).

22. S. J. Eliades, X. Wang, Neural substrates of vocalization feedback monitoring in primate auditory cortex. Nature 453, 1102–1106 (2008).

23. D. M. Schneider, J. Sundararajan, R. Mooney, A cortical filter that learns to suppress the acoustic consequences of movement. Nature 561, 391–395 (2018).

24. D. M. Schneider, A. Nelson, R. Mooney, A synaptic and circuit basis for corollary discharge in the auditory cortex. Nature 513, 189–194 (2014).

25. P. Lakatos, G. Musacchia, M. N. O’Connel, A. Y. Falchier, D. C. Javitt, C. E. Schroeder, The spectrotemporal filter mechanism of auditory selective attention. Neuron 77, 750–761 (2013).

26. P. Lakatos, G. Karmos, A. D. Mehta, I. Ulbert, C. E. Schroeder, Entrainment of neuronal oscillations as a mechanism of attentional selection. Science 320, 110–113 (2008).

27. M. N. O’Connell, A. Barczak, C. E. Schroeder, P. Lakatos, Layer specific sharpening of frequency tuning by selective attention in primary auditory cortex. J Neurosci 34, 16496–16508 (2014).

28. G. Dirlich, L. Vogl, M. Plaschke, F. Strian, Cardiac field effects on the EEG. Electroencephalogr Clin Neurophysiol 102, 307–315 (1997).

29. N. Ille, P. Berg, M. Scherg, Artifact correction of the ongoing EEG using spatial filters based on artifact and brain signal topographies. J Clin Neurophysiol 19, 113–124 (2002).

30. J. A. Jiang, C. F. Chao, M. J. Chiu, R. G. Lee, C. L. Tseng, R. Lin, An automatic analysis method for detecting and eliminating ECG artifacts in EEG. Comput Biol Med 37, 1660–1671 (2007).

31. P. Lakatos, J. Gross, G. Thut, A New Unifying Account of the Roles of Neuronal Entrainment. Curr Biol 29, R890–R905 (2019).

32. C. J. Stam, G. Nolte, A. Daffertshofer, Phase lag index: assessment of functional connectivity from multi channel EEG and MEG with diminished bias from common sources. Hum Brain Mapp 28, 1178–1193 (2007).

33. D. Haslacher et al., In vivo phase-dependent enhancement and suppression of human brain oscillations by transcranial alternating current stimulation (tACS). Neuroimage 275, 120187 (2023).

34. S. A. Saxon, Detection of near threshold signals during four phases of cardiac cycle. Ala J Med Sci 7, 427–430 (1970).

35. C. A. Sandman, T. R. McCanne, D. N. Kaiser, B. Diamond, Heart rate and cardiac phase influences on visual perception. J Comp Physiol Psychol 91, 189–202 (1977).

36. B. B. Walker, C. A. Sandman, Visual evoked potentials change as heart rate and carotid pressure change. Psychophysiology 19, 520–527 (1982).

37. C. A. Sandman, Augmentation of the auditory event related potentials of the brain during diastole. Int J Psychophysiol 2, 111–119 (1984).

38. P. Motyka, M. Grund, N. Forschack, E. Al, A. Villringer, M. Gaebler, Interactions between cardiac activity and conscious somatosensory perception. Psychophysiology 56, e13424 (2019).

39. L. Edwards, C. Ring, D. McIntyre, J. B. Winer, U. Martin, Sensory detection thresholds are modulated across the cardiac cycle: evidence that cutaneous sensibility is greatest for systolic stimulation. Psychophysiology 46, 252–256 (2009).

40. E. Al et al., Heart-brain interactions shape somatosensory perception and evoked potentials. Proc Natl Acad Sci U S A 117, 10575–10584 (2020).

41. E. Al, F. Iliopoulos, V. V. Nikulin, A. Villringer, Heartbeat and somatosensory perception. Neuroimage 238, 118247 (2021).

42. T. Engelen, M. Solca, C. Tallon-Baudry, Interoceptive rhythms in the brain. Nat Neurosci 26, 1670–1684 (2023).

43. L. F. Barrett, W. K. Simmons, Interoceptive predictions in the brain. Nat Rev Neurosci 16, 419–429 (2015).

44. X. Gu, P. R. Hof, K. J. Friston, J. Fan, Anterior insular cortex and emotional awareness. J Comp Neurol 521, 3371–3388 (2013).

45. H. Y. Weng, J. L. Feldman, L. Leggio, V. Napadow, J. Park, C. J. Price, Interventions and Manipulations of Interoception. Trends Neurosci 44, 52–62 (2021).

46. J. Dewey, The theory of emotion: I: Emotional attitudes. Psychological review 1, 553 (1894).

47. W. B. Cannon, The James-Lange theory of emotions: a critical examination and an alternative theory. By Walter B. Cannon, 1927. Am J Psychol 100, 567–586 (1987).

48. S. Schachter, J. E. Singer, Cognitive, social, and physiological determinants of emotional state. Psychol Rev 69, 379–399 (1962).

49. S. S. Khalsa et al., Interoception and Mental Health: A Roadmap. Biol Psychiatry Cogn Neurosci Neuroimaging 3, 501–513 (2018).

50. M. A. Farmer, M. N. Baliki, A. V. Apkarian, A dynamic network perspective of chronic pain. Neurosci Lett 520, 197–203 (2012).

51. E. A. Mayer, Gut feelings: the emerging biology of gut-brain communication. Nat Rev Neurosci 12, 453–466 (2011).

52. J. Radley, D. Morilak, V. Viau, S. Campeau, Chronic stress and brain plasticity: Mechanisms underlying adaptive and maladaptive changes and implications for stress-related CNS disorders. Neurosci Biobehav Rev 58, 79–91 (2015).

53. A. Guerra, V. López-Alonso, B. Cheeran, A. Suppa, Solutions for managing variability in non-invasive brain stimulation studies. Neurosci Lett 719, 133332 (2020).

54. K. Nasr, D. Haslacher, E. Dayan, N. Censor, L. G. Cohen, S. R. Soekadar, Breaking the boundaries of interacting with the human brain using adaptive closed-loop stimulation. Prog Neurobiol 216, 102311 (2022).

55. F. H. Kasten, K. Duecker, M. C. Maack, A. Meiser, C. S. Herrmann, Integrating electric field modeling and neuroimaging to explain inter-individual variability of tACS effects. Nat Commun 10, 5427 (2019).

56. G. Thut et al., Guiding transcranial brain stimulation by EEG/MEG to interact with ongoing brain activity and associated functions: A position paper. Clin Neurophysiol 128, 843–857 (2017).

57. T. O. Bergmann, Brain State-Dependent Brain Stimulation. Front Psychol 9, 2108 (2018).

58. J. Silvanto, N. Muggleton, V. Walsh, State-dependency in brain stimulation studies of perception and cognition. Trends Cogn Sci 12, 447–454 (2008).

59. C. Bradley, A. S. Nydam, P. E. Dux, J. B. Mattingley, State-dependent effects of neural stimulation on brain function and cognition. Nat Rev Neurosci 23, 459–475 (2022).

60. J. S. Brittain, P. Probert-Smith, T. Z. Aziz, P. Brown, Tremor suppression by rhythmic transcranial current stimulation. Curr Biol 23, 436–440 (2013).

61. S. R. Schreglmann et al., Non-invasive suppression of essential tremor via phase-locked disruption of its temporal coherence. Nat Commun 12, 363 (2021).

62. M. Fiene et al., tACS phase-specifically biases brightness perception of flickering light. Brain Stimul 15, 244–253 (2022).

63. M. Fiene, B. C. Schwab, J. Misselhorn, C. S. Herrmann, T. R. Schneider, A. K. Engel, Phase-specific manipulation of rhythmic brain activity by transcranial alternating current stimulation. Brain Stimul 13, 1254–1262 (2020).

64. D. Haslacher, K. Nasr, S. E. Robinson, C. Braun, S. R. Soekadar, Stimulation Artifact Source Separation (SASS) for assessing electric brain oscillations during transcranial alternating current stimulation (tACS). Neuroimage 10.1016/j.neuroimage.2020.117571, 117571 (2021).

65. D. Haslacher, P. Reber, S. Soekadar, Targeting alpha oscillations using closed-loop transcranial alternating current stimulation. Brain Stimulation: Basic, Translational, and Clinical Research in Neuromodulation 16, 235 (2023).

66. B. Zoefel, A. Archer-Boyd, M. H. Davis, Phase Entrainment of Brain Oscillations Causally Modulates Neural Responses to Intelligible Speech. Curr Biol 28, 401–408 e405 (2018).

67. P. C. Miranda, M. Lomarev, M. Hallett, Modeling the current distribution during transcranial direct current stimulation. Clin Neurophysiol 117, 1623–1629 (2006).

68. M. Bortoletto, C. Rodella, R. Salvador, P. C. Miranda, C. Miniussi, Reduced Current Spread by Concentric Electrodes in Transcranial Electrical Stimulation (tES). Brain Stimul 9, 525–528 (2016).

69. N. Grossman et al., Noninvasive Deep Brain Stimulation via Temporally Interfering Electric Fields. Cell 169, 1029-1041.e1016 (2017).

70. I. R. Violante et al., Non-invasive temporal interference electrical stimulation of the human hippocampus. Nat Neurosci 26, 1994–2004 (2023).

71. M. J. Wessel et al., Noninvasive theta-burst stimulation of the human striatum enhances striatal activity and motor skill learning. Nat Neurosci 26, 2005–2016 (2023).

72. K. Nasr, D. Haslacher, S. Soekadar (2023) Towards adaptive deep brain neuromodulation using temporal interference magnetic stimulation. in 10. Transcranial Magnetic Stimulation (TMS) (Elsevier Inc.).

73. R. A. Payne, C. N. Symeonides, D. J. Webb, S. R. Maxwell, Pulse transit time measured from the ECG: an unreliable marker of beat-to-beat blood pressure. J Appl Physiol (1985) 100, 136–141 (2006).

74. A. Gramfort et al., MEG and EEG data analysis with MNE-Python. Front Neurosci 7, 267 (2013).

75. C. R. Harris et al., Array programming with NumPy. Nature 585, 357–362 (2020).

76. P. Virtanen et al., SciPy 1.0: fundamental algorithms for scientific computing in Python. Nat Methods 17, 261–272 (2020).

77. R. J. Croft, R. J. Barry, Removal of ocular artifact from the EEG: a review. Neurophysiol Clin 30, 5–19 (2000).

78. A. Zalesky, A. Fornito, E. T. Bullmore, Network-based statistic: identifying differences in brain networks. Neuroimage 53, 1197–1207 (2010).

79. E. Maris, R. Oostenveld, Nonparametric statistical testing of EEG- and MEG-data. J Neurosci Methods 164, 177–190 (2007).

80. R. VanRullen, How to Evaluate Phase Differences between Trial Groups in Ongoing Electrophysiological Signals. Front Neurosci 10, 426 (2016).

