## Supplementary Materials for "Heartbeat perception is causally linked to frontal delta oscillations"

### Results

#### Frontal delta phase synchrony is not accounted for by evoked brain activity or ocular artifacts

EEG signals exhibited neuronal responses evoked by heartbeats and auditory stimuli, each with rhythmicity in the delta frequency range. A HEP with a maximum positivity approximately 300 ms after the ECG R-peak was observed in parietal and occipital areas (Fig. S1A). An AEP with a maximum positivity approximately 200 ms after the tone onset was observed in frontal midline areas (Fig. S1B). We corrected delta-band EEG signals by removing activity related to HEPs, AEPs, and EOG signals (see methods section "Electroencephalography data processing"). No effect on the spatial pattern of delta phase synchrony due to any of these corrections was observed (Fig. S1C – S1E). Crucially, the difference in frontal delta phase synchrony between incorrect and correct responses in the heartbeat detection task (Fig. 2) was comparable before and after correction for HEPs and AEPs ($t(23) = 0.164$, $p = 0.872$). Likewise, the same difference in frontal delta phase synchrony was comparable before and after correction for ocular artifacts by regression-based removal of EOG signals ($t(23) = -0.516$, $p = 0.611$).

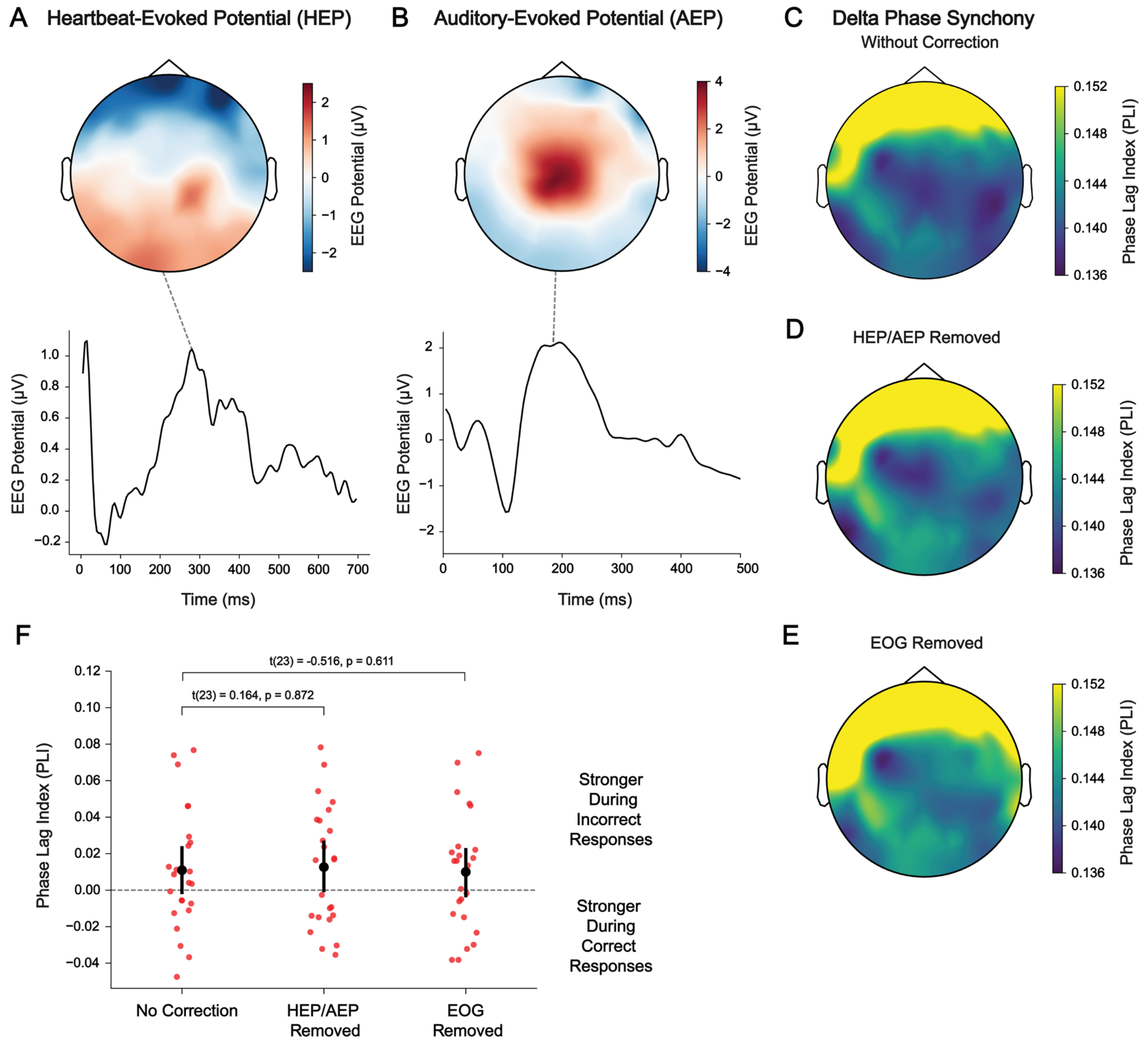


**Figure S1. Frontal delta oscillations are distinct from heartbeat-evoked potentials, auditory evoked potentials, and ocular artifacts. (A)** A heartbeat-evoked potential (HEP) was observed as a positive deflection in parietal and occipital brain areas. **(B)** An auditory-evoked potential (AEP) was observed as a positive deflection in frontal midline brain areas. **(C)** In absence of correction for evoked potentials, delta-band activity exhibited phase synchrony predominantly between frontal brain areas. **(D)** Frontal delta phase synchrony was preserved despite removal of delta-band activity linked to HEPs and AEPs using template subtraction. **(E)** Frontal delta phase synchrony was preserved despite removal of delta-band activity linked to electrooculography (EOG) artifacts using a regression-based approach. **(F)** The difference in frontal delta synchrony between incorrect and correct responses in the heartbeat detection task (Fig. 2) was preserved despite removal of delta-band activity linked to HEPs, AEPs, and EOG artifacts. For details on these signal processing steps, see methods section "Electroencephalography data processing".

### Stimulation artifact source separation recovers frontal delta oscillations targeted by transcranial alternating current stimulation

We first assessed whether delta oscillations could be recovered from EEG data recorded in the presence of AM-tACS by comparing the topography of PLI with that in absence of AM-tACS. Before application of SASS (Fig. S2A), we found that the topography of PLI in the presence of AM-tACS was strongly distorted by the electric stimulation artifact, such that no positive spatial correlation between PLI in absence of AM-tACS and in the presence of AM-tACS could be found ($r = -0.422$, $p = 0.999$). As AM-tACS was applied to the frontal cortex, the PLI attenuated artifactual synchrony in this region, resulting in a dominant contribution of occipital brain areas (Fig. S2B). After application of SASS (Fig. S2C), the spatial pattern of PLI in the presence of AM-tACS was comparable to that in absence of AM-tACS ($r = 0.641$, $p = 5.79\times10^{-5}$). Thus, the expected pattern of frontal delta synchrony was restored by SASS.

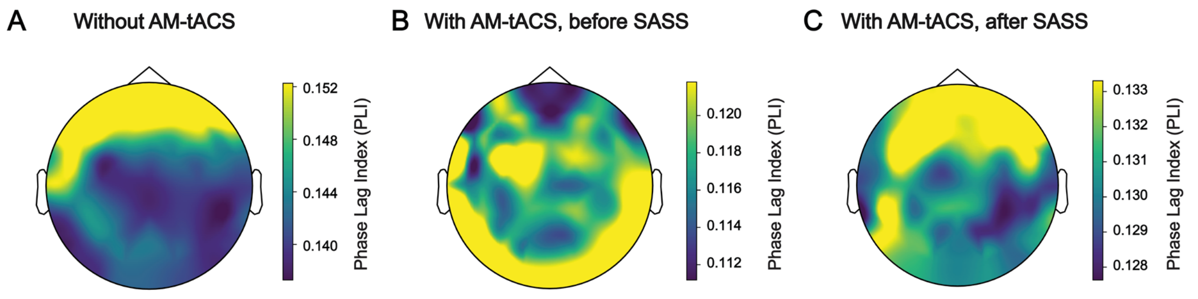


**Figure S2. Stimulation artifact source separation (SASS) recovers delta synchrony in the presence of amplitude-modulated transcranial alternating current stimulation (AM-tACS). (A)** Electroencephalography (EEG) data in absence of AM-tACS revealed prominent delta synchrony in frontal brain regions. **(B)** In the presence of AM-tACS, delta synchrony was obscured by an electric stimulation artifact. **(C)** After application of SASS to the EEG data, delta synchrony in the presence of AM-tACS was comparable to delta synchrony in absence of AM-tACS. The PLI values depicted here reflect the average PLI across all pairwise connections involving each EEG electrode and are averaged across participants.

### Frontal delta phase synchrony was modulated by transcranial alternating current stimulation in a phase-dependent manner

To further assess phase-dependent modulation of frontal delta synchrony (Fig. 3), we first assessed the phase difference between AM-tACS and frontal delta oscillations when AM-tACS was applied early and late relative to the heartbeat. Within participants, we found that the phase difference between AM-tACS and frontal delta oscillations was clustered around opposite values during early versus late AM-tACS (permutation test, $p = 3.52 \times 10^{-4}$). However, the phase difference between AM-tACS and frontal delta oscillations varied across participants in each delay condition (Fig. S3). Furthermore, within participants, we found that frontal delta phase synchrony was modulated by the phase difference between AM-tACS and frontal delta oscillations (permutation test, $p = 3.14 \times 10^{-10}$). However, the optimal phase difference to enhance or suppress frontal delta synchrony varied across participants (Fig. S4).

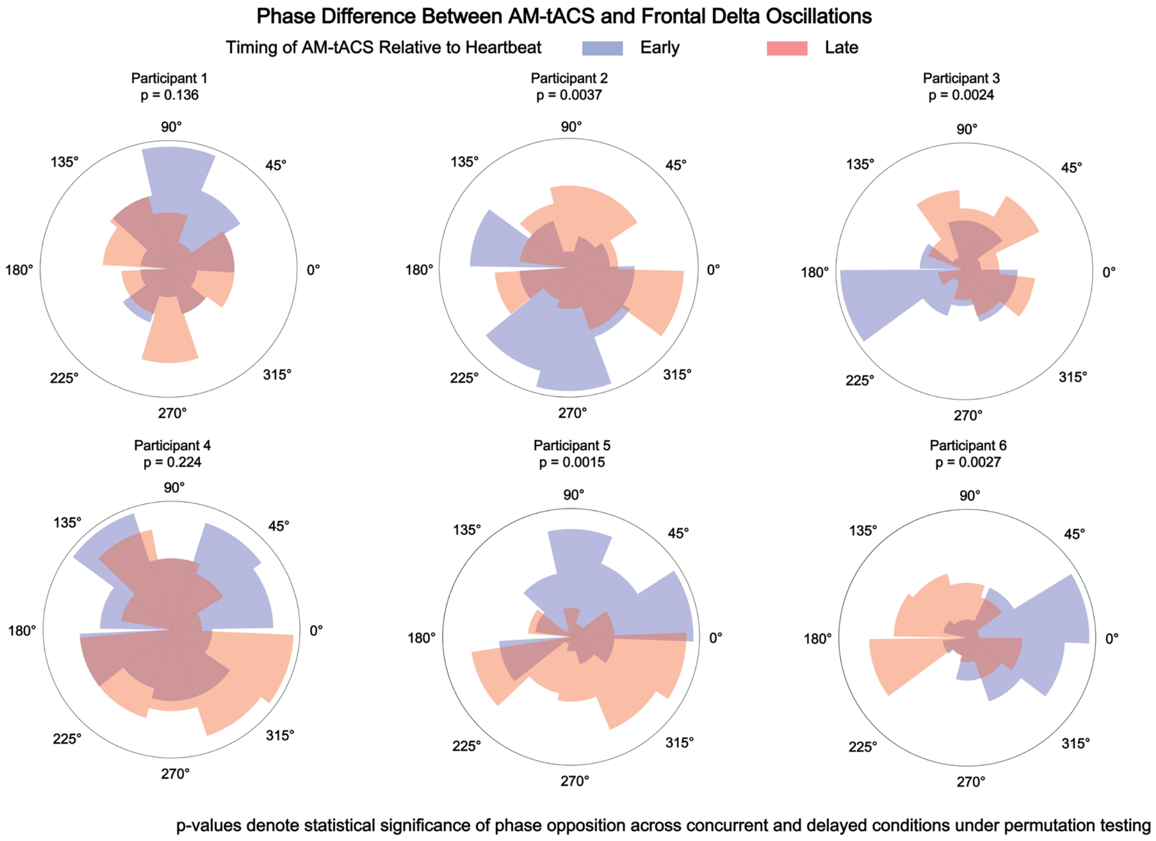


**Figure S3. Amplitude-modulated transcranial alternating current stimulation (AM-tACS) was applied at two opposing phases of frontal delta oscillations.** When evaluating data from each participant individually, the phase difference between AM-tACS and frontal delta oscillations exhibited opposing values during the two AM-tACS delay conditions (see Fig. 1). However, an inter-individual variability in the phase difference between AM-tACS and frontal delta oscillations was observed. Data from six selected participants is depicted.

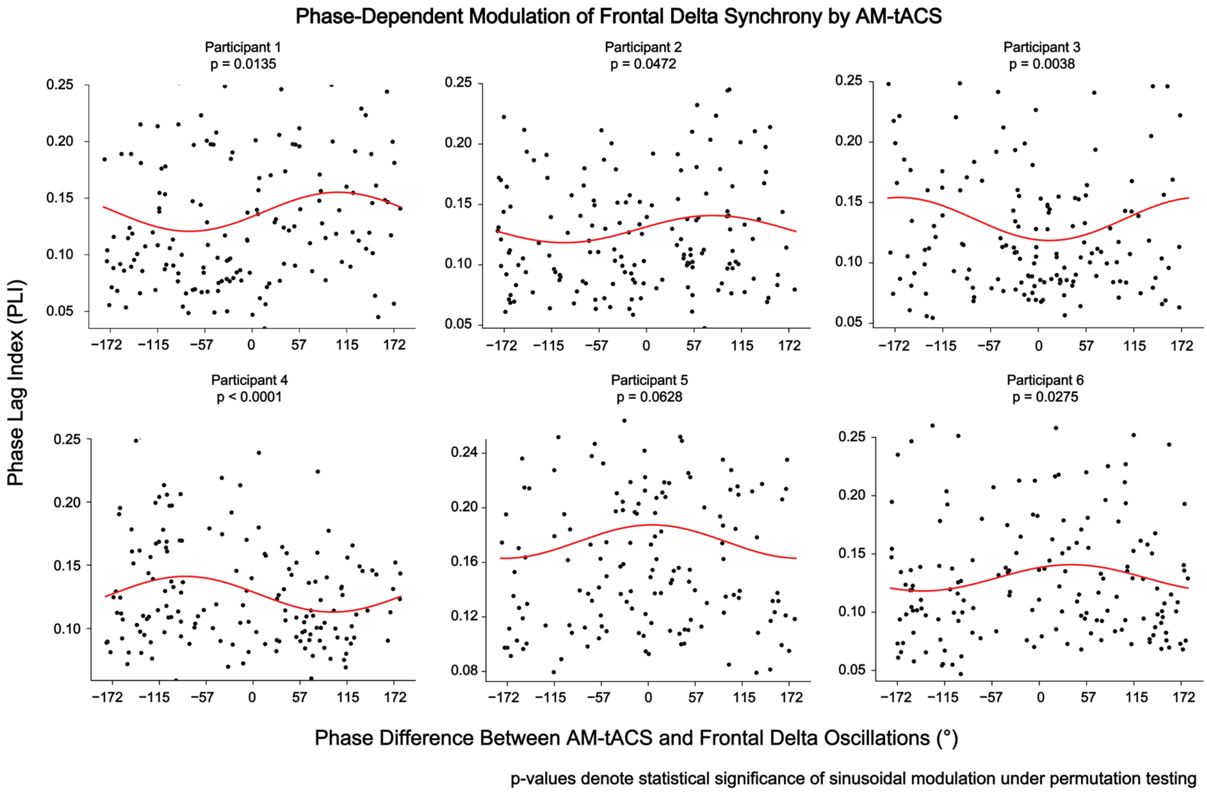


**Figure S4. Frontal delta phase synchrony was modulated by amplitude-modulated transcranial alternating current stimulation (AM-tACS) in a phase-dependent manner.** Assessment of trial-by-trial data for each participant revealed that frontal delta phase synchrony depended on the phase difference between AM-tACS and frontal delta oscillations. However, the optimal phase to enhance or suppress delta synchrony varied across participants. Data from six selected participants is depicted.
